# The effect of information content distributions on word recollection and familiarity

**DOI:** 10.1101/2023.08.02.551617

**Authors:** Joel C. Wallenberg, Salsabila Nadhif Fadhilah, Taylor D. Hinton, Tom V. Smulders, Jenny C.A. Read, Christine Cuskley

## Abstract

This study builds on work on language processing and information theory which suggests that informationally uniform, or smoother, sequences are easier to process than ones in which information arrives in clumps. Because episodic memory is a form of memory in which information is encoded within its surrounding context, we predicted that episodic memory in particular would be sensitive to information distribution. We used the “dual process” theory of recognition memory to separate the episodic memory component (recollection) from the non-episodic component (familiarity) of recognition memory. Though we find a weak effect in the predicted direction, this does not reach statistical significance and so the study does not support the hypothesis. The study does replicate a known effect from the literature where low frequency words are more easily recognized than high frequency ones when participants employ recollection-type memory. We suggest our results may be explained by linguistic processing being particularly adapted to processing linear sequences of information in a way that episodic memory is not. Episodic memory likely evolved to deal with unpredictable, sometimes clumped, information streams.

## Introduction

Information theory [1] characterizes the dynamics of a variety of systems in which information is passed from a sender to a receiver, including the transmission of information across neuronal networks [2] or DNA translation and transcription [3]. Recently, the application of information theory to language has led to the discovery that people tend to distribute information as evenly as possible across utterances as they speak, which helps to maintain effective communication in the presence of “noise” (i.e. interference) of various types [4–7]. This may reflect a general bias for distributing information evenly in transmission in information theoretic systems more generally. For example, abnormally clumped neuronal spikes have been connected to Parkinson’s Disease [8] and chronic tinnitus [9], indicating that clustered information disrupts normal neuronal functioning. In genetics, redundancy spreads information in an adaptive way: there are over three times as many unique mRNA codons as amino acids they code for. Because the same amino acid is coded for by multiple similar codons, it is less likely that a single nucleotide mutation alters the amino acid than if each amino acid had a unique code [10].

Memory encoding can also be viewed in information theoretic terms, as the process of information transfer from the outside world to brain circuits. If the same principles apply to this information transfer as those found in language, and suggested for neural systems and DNA, then clumps of information should inhibit encoding. In other words, more information should be encoded when the incoming information is distributed more evenly in temporal terms. The memory system that is dedicated to storing and retrieving unique events in context (such as unique sequences of words) is called episodic memory [11, 12]. Here, we build on the “‘dual process” theory of recognition memory. [13–15] Familiarity is the recognition that a particular item has been encountered before, but without retrieval of the surrounding context (i.e., whether or not you believe you have seen an item). Recollection, on the other hand, is understood as recognition of an item or word *and the episodic context in which it was encountered*. The two processes are hypothesized to be separable both cognitively and neurobiologically [14, 16–18]. One way to separate the two processes behaviourally is by the use of Receiver Operating Characteristic (ROC) curves, which are fit for each participant so as to calculate the contributions of recollection and familiarity by estimating the ROC’s y-intercept parameter (for recollection) and curvature parameter (“*d*′”, for familiarity [19]). To do this, participants are first presented with a target word list during a study phase. After a retention interval, they are then presented with another list of words, made up of the old words combined with distractor words. Participants are then asked to identify which words they had seen before, and to report their confidence for each judgement. The confidence measures are used to fit the ROC curve.

To investigate how the episodic memory system deals with information that is distributed in different manners throughout a temporal stream, we added a manipulation of the word order in the study phase, so that some participants saw all low frequency words (i.e. high information content words) and high frequency words (i.e. low information content) sorted in ascending or descending order (“Clumped”), while others saw a list of words in which low and high frequency words alternated (“Even”). The Even order, which spreads information across the sequence as a whole in a more distributed fashion, is known to be resistant to catastrophic effects of certain types of noise [20, 21]. We therefore hypothesize that episodic memory encoding will be more successful when participants see sequences where information is more evenly distributed, *when it is a sequence that is being stored*. In other words, we predict this effect for recollection type memory *only*, which stores and retrieves a target item with its surrounding context.

## Materials and methods

### Participants

#### Sample A

The experiment was approved by the Newcastle University Ethics Committee, REF 232/2020, and all research was performed in accordance with relevant guidelines and regulations. Data collection for this sample took place between 10 February 2020 and 20 July 2020. Written informed consent was obtained from all participants prior to the start of the experimental task. A total of 315 participants were recruited on Amazon Mechanical Turk (paid $4 for completing the task) and 10 participants volunteered on social media. The task took approximately 20-30 minutes in piloting. Participants took on average 22 minutes to complete (SD= 12 minutes); some participants completed extremely quickly (e.g., within 7 minutes) while others took over an hour. Due to the unmonitored online nature of the task, participants with completion times shorter than 1 standard deviation below the mean completion time and greater than 2 standard deviations longer than the mean were excluded, leaving 284 participants (186 male, 98 female, 1 preferred not to say) for analysis. Only two volunteers were eliminated and the remainder were Mechanical Turk workers (of 285 participants analysed, 277 were recruited via Mechanical Turk and 8 volunteered via social media). Participants included in analyses had ages ranging from 19 to 72 years old (mean ≈ 37.3, *SD ±* 11.6), and completion times in the range of 10-51 minutes, with a mean completion time of 21.55 minutes.

139 participants were in the smooth condition (shown a list of words with alternating high and low information content, making for a smoother distribution of information across the list), and 147 were in the clumped condition (shown a list of words sorted by information content, such that information is clumped within the list).

#### Sample B

The experiment was approved by the Newcastle University Ethics Committee on 18 March 2021, and all research was performed in accordance with relevant guidelines and regulations. Written informed consent was obtained from all participants prior to the start of the experimental task. Data collection for this sample took place between 5 April 2021 and 12 July 2021. A total of 199 participants were recruited on Amazon Mechanical Turk. For this sample, the word memory/recollection task occurred alongside an image memory/recollection task not analysed here: participants either randomly completed the image portion of the task or the word portion of the task first. Due to the variable nature of completion times in the first task, this task was time limited: participants had to submit the completed task on Mechanical Turk within one hour of accepting it. Given the longer overall nature of this task and the time-limited completion, participants were paid $12 for completing this task (all participants who completed the task in Sample A were excluded based on their Mechanical Turk Worker ID). Completion times for the word block of the task on its own were not collected; however, participants were excluded from analyses based on their overall completion time as in Sample A. Here, the mean completion time was 29 minutes, with a minimum of 14 minutes and a maximum of 59 minutes (as this was the maximum permitted by the task), with an standard deviation of 7 minutes. As with Sample A, participants with completion times less than 1 SD below the mean (22 minutes) and more than 2 SDs above the mean (43 minutes) were excluded.

This left a total of 173 participants included in analysis. 87 were in the smooth condition, and 86 were in the clumped condition.

### Materials

A total of 140 words were chosen from the English Lexicon Project [22], including 70 target words and 70 distractors. All words were two syllable monomorphemic nouns between 5-8 characters, and fell into three frequency categories based on their log frequency in the HAL corpus [23]. Of the 70 target words, 35 were low frequency (log frequency between 10.12 and 12.55) and 35 were high frequency (log frequency between 2.57 and 6.68). The 70 distractor words were of a mid range; log frequency between 7.16 and 9.89 (overlapping with neither the high or low categories). Materials were identical for both samples.

The target word lists were presented in four potential orders, according to the smooth vs clumped condition, and whether a list started with high frequency words or low frequency words (low start and high start respectively). In lists which had clumped information distributions, words were sorted either in ascending (from low to high) or descending (from high to low) order according to their frequency. In conditions which had even information distributions, the words in each frequency grouping were sorted in ascending order separately, and then alternately appended to a central list pulling from the start of one list and the end of the other. In other words, for the high start/even condition, the low frequency and high frequency words were each arranged in ascending order, and then rearranged into a central list which was assembled by removing the first word from the high frequency list and the last word from the low frequency list iteratively until both the original lists were empty. Participants were systematically assigned a condition as described below (in “Procedure”). The list of words for rating (including all low frequency, high frequency, and mid frequency words for a total of 140 items) was presented in a random order for each participant (using the random.shuffle() function in Python).

### Procedure

The task was conducted in the browser using JavaScript and jQuery, with a Python/Flask server deployed on Heroku (https://heroku.com), and data stored in MongoDB Atlas. Fully documented code for the experimental setup is available at https://github.com/CCuskley/Wordmemory/tree/main/Materials, and can be demoed at https://exps-main.herokuapp.com/wmdemo (Note that this demo only shows the low start smooth list of words invariably, although the open source code includes condition assignment.)

Each target word was shown in the centre of a white screen, and target words alternated with a fixation cross. Each target word displayed for 1000ms and the fixation cross displayed for 500ms between words.

Participants were randomly allocated to a condition based on how many previous participants had completed the task in that condition. Participants began the experiment by consenting to standard terms of participation, including details about compensation, length of the task, and data use. They were then given very brief instructions indicating approximately how long the experiment would take, and explaining that they would be tasked with recalling a list of words. Before proceeding to the main task, participants were first given a simple test to prevent bots or scripts from completing the task. If they failed this task, they could not view or proceed to the remainder of the experiment, and thus could not complete the task, either as a volunteer or on Mechanical Turk.

After passing this test, participants were asked to provide their age and gender (male, female, other, or prefer not to say). They were then asked whether their first language is English. If they said yes, they were asked if they knew other languages. If they said ‘no’, they were asked for the age at which they started learning English, providing us with a rough proxy for proficiency when combined with their current age.

Following this, they were given more detailed instructions, that they would see a list of words, watch a video, and then try to recall words from the list. First, participants were shown a short demo list to become familiar with how the list would be displayed for the target words; this simply displayed each word in the sentence ‘This is where the words will show try to remember them’ in succession. None of the words in this sentence were part of the target or distractor set used in the main task. Before moving onto the target list, participants were advised that the target list would include unrelated words, and initiated the list of 70 target words. After this, participants watched a short (3min) video (with no lyrics or narration) of either cats (https://vimeo.com/212247939 or plants (https://vimeo.com/69225705), before being given brief instructions on the recall phase. In the recall phase, they were shown all 140 words (the 70 targets and 70 medium frequency distractors) in a random order. Each word was shown in isolation with the questions “Did you see it?” (answer was yes or no button) and “How sure are you” (answers were “Not at all”, “Sort of” and “Very”). Participants could change their answers before moving onto the next word, and a progress bar at the top of the screen displayed how far along they were. When they had completed the entire list the task ended.

## Analysis

### Fitting

The fitting of Receiver Operating Characteristics for each participant, and in the second analysis for high frequency words and low frequency words for each participant, was conducted with purpose-built scripts in R, which can be found in the Github repository below. All other statistical analysis used R (core libraries, [24] *lme4* for mixed-effects models, [25] and *ggplot2* for plots [26]).

#### Mathematical model

The model assumes that during the recall phase, each word elicits an internal signal indicating its familiarity. This signal is assumed to be normally distributed and to have unit variance. The mean familiarity is *d*′ for words that were in the previously-seen target list, and 0 for words that were not. Words that were in the target list may also be recollected via episodic memory, with probability *R*. We assume that participants judge a word as “previously seen” if the familiarity signal exceeds a criterion value *C*, or (for words in the target list) if they recollect it. An observer’s performance is therefore characterized by the three parameters *d*′, *R* and *C*. Where the word was in the target list, the probability that the observer correctly identifies it as such is

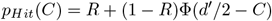

where Φ is the cumulative distribution function of the standard normal distribution. The probability they class it as not seen is

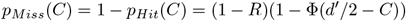

For words not in the target list, the probability that the observer makes a “false alarm” is

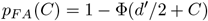

while the probability that they correctly reject it is

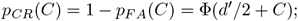

#### Estimating the ROC curve

We have written these as functions of the decision criterion *C* because we assume that, whereas *R* and *d*′ are fixed for a given observer and experimental condition, *C* can vary. The Receiver Operating Characteristic curve, or ROC curve, is obtained by plotting *p*_*Hit*_(*C*) against *p*_*F A*_(*C*) as *C* varies.

We estimate the effect of varying *C* by rescoring observers based on their stated confidence. We simulate a high decision criterion by recoding as “no” trials where observers actually responded “yes” the word was in the target list but indicated that they were “not at all” or “sort of” sure; only trials where observers said “yes, very” were retained as “yes”. This reduces both the probability of hits and the probability of false alarms. We can then lower the decision criterion by recoding as “no” only trials where observers answered “yes” but were “not at all sure”, and so on. In this way, we obtain estimates of 5 points on the ROC curve. We use this to fit maximum-likelihood estimates of *R* and *d*′, the parameters of interest. This also involves fitting 5 nuisance parameters, the decision criteria, *C*_*j*_.

#### Fitting procedure

For a given *R,d*′ and *C*, corresponding to a single point on an ROC curve, the log-likelihood of getting a particular set of results is

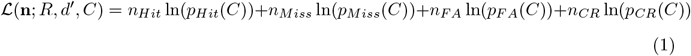

where the vector **n** represents the number of trials in each of the four categories. For given values of *R, d*′, we first optimize *C*_*j*_ individually for each of the 5 confidence boundaries **n**_*j*_:

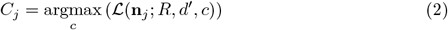

This corresponds to fixing the ROC curve, and sliding points along it to match the data. Then we seek the values of *R* and *d*′ which maximize

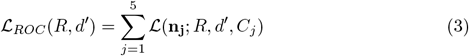

subject to the bounds *R* = [0, 1) and *d*′ = [0, 2]. The upper bound of *d*′ is for practical reasons: very large *d*′ cause numerical overflow and are not needed to model the data anyway. We bound *d*′ at 0 because negative values correspond to performance below chance.

#### 0.0.1 Optimization starting point

In optimization problems, a suitable initial guess is often critical. To obtain this, we first look at the proportion of correct rejections in the observer’s actual judgments (i.e. without any recoding), *P*_*CR*_. From the equations above, we expect 0.5*d*′ + *C* = Φ^−1^(*P*_*CR*_). Since *d*′ is bounded at 0, if Φ^−1^(*P*_*CR*_) *<* 0, we set *d*′ = 0 and *C* = Φ^−1^(*P*_*CR*_); otherwise, we set *C* = 0 and 0.5*d*′ = Φ^−1^(*P*_*CR*_). We can then estimate R from

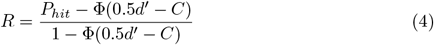

These values are taken as the starting-point for the R optimization routine “optim”, using method “L-BFGS-B” with lower, upper bounds set to 0, 0.999 for *R* and 0, 2 for *d*′. We also checked that the same results, to within the tolerance, were obtained with the MATLAB function “fminsearch”, with the cost function set to infinity when *R <* 0 or *d*′ *<* 0.

#### Confidence intervals

We estimate confidence intervals on *R* and *d*′ using the likelihood ratio approach. The 95% confidence intervals correspond to the contours *ℒ*_95_ = *ℒ*_*max*_ − 1.92. Having obtained the values *R*_*fit*_ and *d*′_*fit*_ which give the maximum log-likelihood *ℒ*_*max*_, for each *R* we seek the maximum and minimum *d*′ for which *ℒ*_*ROC*_(*R, d*′) *> ℒ*_95_. The maximum and minimum values encountered over all *R* are taken as the 95% confidence interval on *d*′. The 95% confidence interval on *R* is obtained similarly.

#### Statistical analysis

Distributions of all three fitted ROC parameters were highly skewed, with a peak at zero (cf Fig 1, Fig 2), so are very poorly approximated by a normal distribution. Accordingly, we fitted the parameters with a gamma distribution using a log link function. To avoid numerical problems with very small values and to avoid taking the log of zeros, all fitted values of *R* and *d*′ ¡0.01 were set equal to 0.01 for analysis and display. The gamma distribution has two parameters: a shape parameter allowing for different amounts of skew, and a scale or rate parameter controlling the variance. In fitting the models, the shape parameter was assumed to be the same for all participants and conditions, while the scale parameter was allowed to vary. Mixed models were fitted using function glmer from R package lme4.

**Fig 1.**
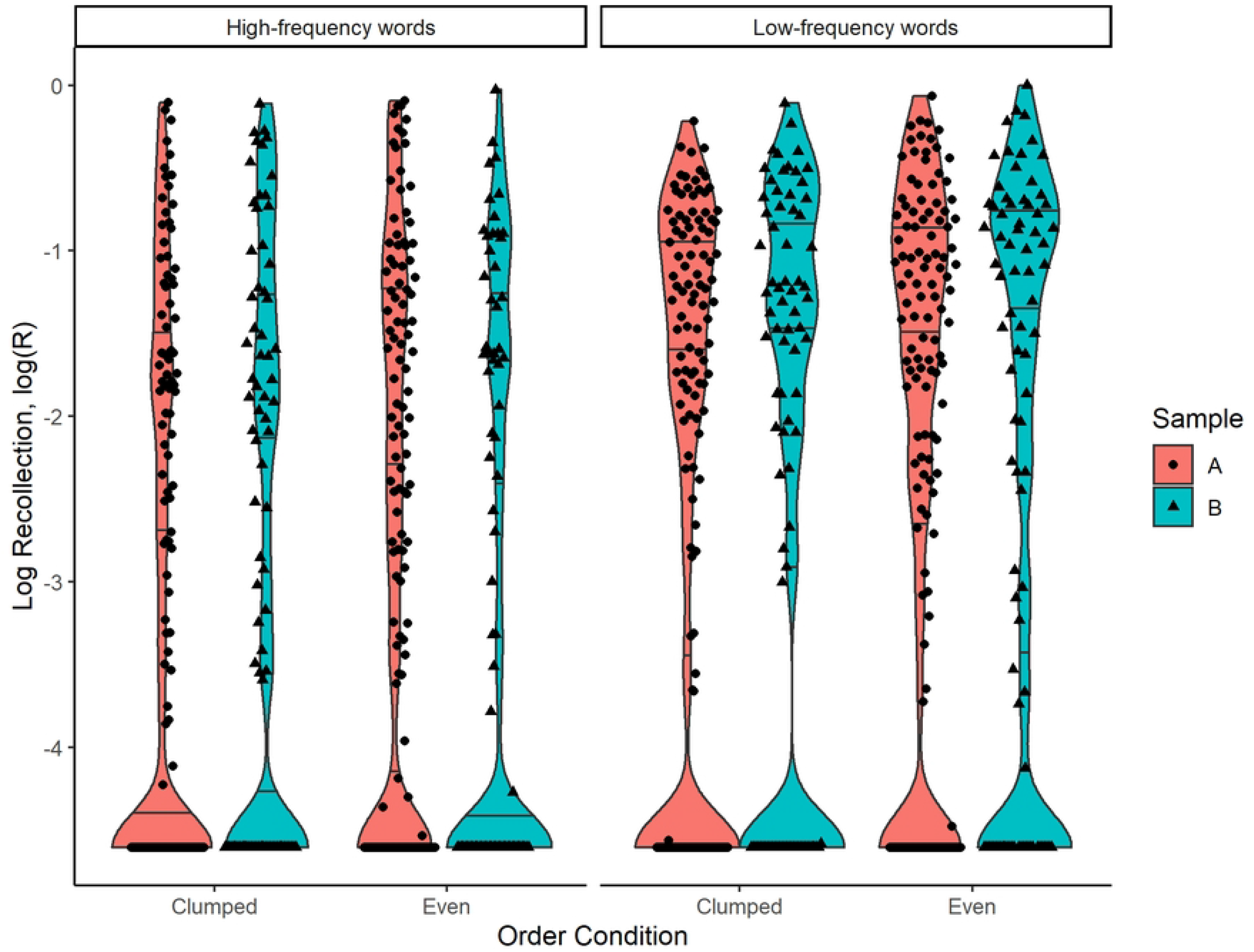
Distribution of logarithm of fitted recollection parameter log(R), fitted separately to high-frequency and low-frequency words. Distributions are shown separately for Clumped and Even conditions, and for the two samples (see Methods). The three lines on each distribution mark the 25%, 50% and 75% quantiles. The large number of points at log(*R*)=-4.6 represents all fitted values ¡0.01, which were set to a nominal value of 0.01 for analysis and display.

**Fig 2.**
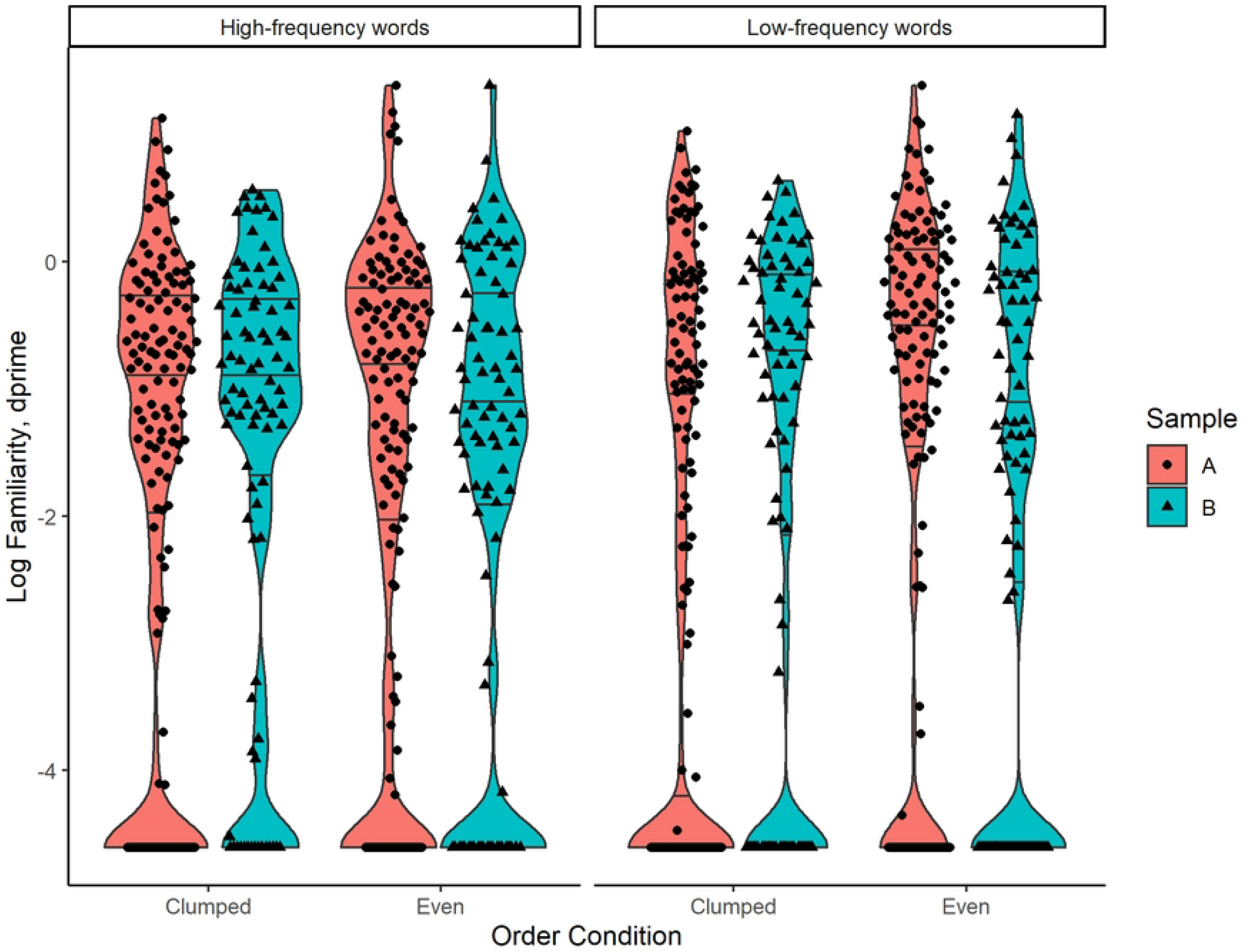
Distribution of logarithm of fitted familiarity parameter dprime, fitted separately to high-frequency and low-frequency words. Other details as in Fig 1.

## Availability of data and materials

The datasets generated and analyzed during the current study are available in the Wordmemory/DataAndAnalyses/ repository, https://github.com/CCuskley/Wordmemory

## Results

### Unusual words are recollected better

Previous studies have found some significant, though conflicting, effects of word frequency on the recollection of individual words [14] (esp. pp.466–467). In order to distinguish the effect of word frequency *per se* from the distribution of information across the sequence, we first split each participant’s data into scores for low frequency words and scores for high frequency words, and fitted ROC curves separately to each subset. We thus have two observations per participant for each parameter (i.e. the parameter calculated only on low frequency words, and the parameter calculated only on high frequency words).

Note that we performed the recollection experiment twice, in two different years, and will refer to the results of those experimental runs as “Sample A” and “Sample B” (see Methods for details). Results are discussed with respect to both samples unless otherwise indicated (with effect of “Sample” controlled for statistically).

Overall performance was significantly better for the low-frequency words (*p <* 10^−6^, *β* = 0.040, mixed effects gamma regression of AUROC on frequency). This was because the low-frequency words were recollected better (*p <* 10^−15^, *β* = 0.63, ROC *y*-intercept *R*), and not because they were more familiar on re-presentation (*p* = 0.27, *β* = −0.082, ROC *d*′). The effect on recollection is illustrated in Fig 1, where it is clear that *R* is consistently much higher for the low-frequency words. Fig 2 shows the lack of an effect for dprime.

### Information distribution did not have a detectable effect

Fig 1 and Fig 2 show summary statistics for the conditions where high and low frequency words were evenly distributed (*Even*) vs where they were ordered (*Clumped*). While in both groups *R* is a little higher for the Even condition, this difference is not significant. In a mixed effects gamma regression of *R* on frequency and order, the effect of frequency was unchanged (*p <* 10^−15^, *β* = 0.63) while the effect of order was not significant (*p* = 0.40, *β* = 0.12). The AIC and BIC were both higher (i.e. worse) for the model including order as well as word frequency. There was also no significant interaction with sample.

Because we halve the number of trials per participant when we fit separately to high and low frequency words, the estimates of ROC parameters are noisier. To investigate whether this was obscuring the effect of information distribution, we also examined fitting all words for each participant with a single ROC curve.

There was now a hint of an effect of order on recollection (*p* = 0.078, *β* = 0.17, fixed-effect gamma regression of *R* on order), though not on overall performance (*p* = 0.15, *β* = 0.026, AUROC) nor on familiarity (*p* = 0.21, *β* = 0.13, *d*′).

The effect of order was significant in Sample A (*p* = 0.01, *β* = 0.31, fixed-effect gamma regression of *R* on order), but not in Sample B (*p* = 0.77, *β* = −0.05). However, the AIC and BIC are both higher (worse) for a model including sample as a regression parameter, suggesting that there is not a genuine difference between samples. Overall, therefore, we do not have evidence for an effect of order.

We also examined whether time taken to complete the task correlated with performance. Very low or very high completion times were excluded from analyses, and participants in Sample A had a wider range of completion times than in Sample B due to task differences (see Methods for details). Overall, the 9/284 participants from Sample A who spent more than 45 minutes on the task performed slightly better. Once these participants were removed, there was no relationship between time taken and performance (*p*=0.19, *β* = 0.0014 per minute, fixed-effect gamma regression of AUROC on time taken in minutes).

## Discussion

The results did not support our hypothesis that recollection memory benefits from receiving stimuli in an informationally even order rather than a more clumped order. We believe this points to an important difference between the word recognition task and linguistic processing in ecologically valid conditions.

Literature on information theory and language has shown that, given the choice, speakers tend to produce utterances which avoid major peaks in their information content, possibly because such “even” distributions mitigate against information loss due to noise events [4, 5, 20, 27]. Sequences which distribute information more uniformly also tend to lose information to noise in uniform amounts, avoiding the potentiality for catastrophic noise events, e.g. where a majority of a sequence’s information is destroyed [21]. Such catastrophic noise events run the risk of making an entire sequence of information-bearing symbols useless to a receiver, assuming the receiver cares about interpreting or processing the sequence as a whole, i.e. the symbols are evaluated with respect to each other in some way (as words are in a sentence).

Our hypothesis was based on the idea that the episodic memory might encode memories as sequences of symbols, particularly if people were presented with stimuli to remember in a sequence. Under the dual process model of episodic memory, recollection involves the retrieval of a recognized item with the context in which the item was encountered [15, 17–19]. If the target item and its context were stored and retrieved as an ordered sequence of symbols (or a few such sequences), then we might reasonably expect recollection to function under similar constraints to the linguistic processing of a sentence. Thus, recollection might similarly prefer sequences that are more even in their information distributions, and so more robust to noise events. (Note that we did not expect familiarity to process items in sequences, since familiarity does not involve retrieval of an item’s context in the dual process model.) In our experiment, the items to be remembered were temporally ordered words in a list, and so the items in their context in this case were particularly amenable to storage and retrieval as ordered sequences. The words crucially did not form phrases or sentences, so that we could see if recollection preferred informationally even sequences when there was no sentence-processing task at play.

Our results show no evidence that recollection does prefer even sequences, however, so recollection may operate quite differently from linguistic processing, and perhaps does not process memories in sequences at all. This null result is perhaps unsurprising for a number of reasons. First, the task of linguistic processing always involves a temporally ordered sequence of symbols, so the language processing system will be naturally adapted to preserving information in temporal sequences of symbols. Even if some parts of the linguistic signal are processed in parallel (e.g. intonation and word identity in spoken languages, or hand shape and hand movement in signed languages), there is still always some important temporal ordering of linguistic symbols to be processed. Additionally, linguistic processing deals not just with sequences of symbols, but with meaningful sequences of symbols: the ordering of sounds phonemes, morphemes, words, phrases and their composition into larger groupings (constituents) is itself meaningful, and so crucial to linguistic communication. General episodic memory, on the other hand, may well not be adapted to processing ordered lists of items. Just as language processing is adapted to temporally ordered elements, episodic memory will be adapted to the way people generally encounter the variety of meaningful stimuli that make up episodes of their experience. Much of this information will be visual, and so ordered in multiple dimensions with no one dimension privileged, and there will be overlapping stimuli from other senses making up each episode that recollection processes. Such layered and multi-dimensional input could theoretically be digitized into an ordered list brain-internally, just as photographs can be digitized into binary sequences, but it also may not be (or not at a level we can detect with this kind of experiment).

Our results also show that word frequency was a good predictor of recollection performance, with lower frequency words (i.e. higher information content) being recognized more accurately. There was no significant effect of word frequency on the *d’* familiarity parameter in our sample. The increase in recollection with low frequency words and the lack of an effect on familiarity are both consistent with the majority of previous studies (e.g. [28], review in [14], discussion of the *word frequency mirror pattern* in [29]) though a few report recollection effects in the opposite direction. [14] There is also some evidence that frequency can affect familiarity performance in different directions depending on the recency of stimuli. [29] The recollection effect may in part be a general attention or surprise effect, eliciting a variety of brain and sympathetic nervous system responses that are known to be associated with novel, unfamiliar, surprising, or contextually deviant stimuli [30]. Schomaker & Meeter (2015) also review a body of findings suggesting that these responses to unexpected stimuli may aid the memory encoding of such stimuli, and confer attentional advantages, e.g. in improving focus on a specific task (at least in the short term) [30]. To the extent that the low frequency words in our experiment were generally novel or unexpected to the participants, we might expect novelty responses to translate into main effects of word frequency on overall memory performance.

The greatest limitation of the study was surely that participant recruitment and data collection was all conducted online, due to restrictions stemming from the global pandemic. Researchers could therefore not control the recruiting pool very tightly, nor could we control the setting in which participants completed the experiment, or monitor participants’ behaviour during experimentation. We know from time-to-completion data that some participants completed the task far too quickly for their results to be treated as reliable. Others took an unusually long time to complete the task, and our results show that their overall performance was better when they took longer than 45 minutes. These participants could conceivably have taken notes or otherwise introduced noise into their performance. We excluded both of these types of participants post hoc, but we do not know what else participants might have done e.g. in their own homes, or how their attention might have wandered in various settings. The fact that we found a possibly borderline effect of stimulus order on recollection when we calculated parameters on the maximum number of data points per participant suggests that it would be worth replicating these results in a laboratory environment. Another limitation and direction for future research is the recognition task itself, which is known throughout the dual process literature to involve both recollection and familiarity. This was a reasonable choice, as we did predict a contrast between recollection and familiarity. However, a future study could introduce the “clumped” vs “even” manipulation to a paradigm which specifically targets recollection [18].

## Conclusions

The present study is the first attempt to test whether the information distribution of stimuli has an effect on episodic memory encoding and retrieval, as has been argued for human language processing. We did not observe such an effect, which may point to different evolutionary trajectories for the general episodic memory system and linguistic processing. The latter may well be specifically adapted to preserving information in meaningful sequences of symbols. Episodic memory, however, may not benefit from such a specific adaptation because of the much wider array of stimulus configurations it must process in the natural environment. In addition to this null result, our study replicated a known result: recollection memory performs better on low frequency words than on high frequency words.

## Acknowledgments

We would like to thank members of the Centre for Behaviour and Evolution at Newcastle University and the Experimental Linguistics Lab at the University of York for helpful discussion at presentations of preliminary versions of this material. We would also like to acknowledge that parts of this work were funded by the Economic and Social Research Council (ESRC, UK) Secondary Data Analysis Initiative grant #ES/T005955/1.

## CRediT Author contributions statement

S.F.: Investigation, Formal analysis, Project administration; T.D.H.: Investigation, Formal analysis, project administration, Writing-Review and editing; J.C.W: Conceptualization, Methodology, Formal analysis, Writing-Original draft, Supervision, Project administration, C.C.:Conceptualization, Methodology, Software, Data curation, Writing-Review and editing, Supervision, Project administration; T.V.S.:Conceptualization, Writing-Review and editing, Supervision ; J.C.A.R Software, Formal analysis, Writing-Review and editing, Visualization.

